# The cost of cognition: Measuring the energy consumption of non-equilibrium computation

**DOI:** 10.1101/2025.06.18.660368

**Authors:** Gustavo Deco, Yonatan Sanz Perl, Andrea Luppi, Silvia Gini, Alessandro Gozzi, Shamil Chandaria, Morten L. Kringelbach

## Abstract

In biological systems, survival is predicated on an animal being able to perform computations quickly on a minimal energy budget. What is the energy consumption of non-equilibrium brain computation, i.e., what is the cost of cognition? Previous literature has estimated the metabolic cost using neuroimaging measures of glucose consumption but complementary to these findings, here we directly estimate the computational costs by combining the new field of stochastic thermodynamics with whole-brain modelling. We developed the COCO (COst of COgnition) framework using an analytical expression quantifying the links between energy cost, non-equilibrium and information processing for any given brain state measured with neuroimaging. Importantly, this key relationship also holds at the level of individual brain regions. We used this to quantify the benefits of information processing on the highly anatomically, interconnected hierarchical systems of the brain. Crucially, in empirical neuroimaging data we demonstrate that the human brain uses significantly less energy overall than other mammals (including non-human primates and mice), suggestive of an evolutionary optimisation of the effectiveness of computation. Focusing on the cost of cognition, using large-scale human neuroimaging data of 970 healthy human participants, we show that the resting state uses significantly less energy that seven different cognitive tasks. Furthermore, different kinds of tasks require different amounts of non-equilibrium, information processing and energy consumption. We found that tasks requiring more distributed computation also use more energy. Overall, these results directly quantify the cost of cognition, i.e., the non-equilibrium and energetic demands of information processing, allowing a deeper understanding of how the brain compute in a way that is far more energy efficient than current generations of digital computers and artificial intelligence.

## Introduction

Survival and reproduction are the main aims of biological evolution and reducing energy requirements is a central concern. Any living organism is constantly exchanging a flux of matter and energy with its environment. The Nobel Laurate Ernst Schrödinger proposed that sustaining life is exactly predicated on avoiding equilibrium: “How does the living organism avoid decay? … By eating, drinking, breathing and … assimilating. The technical term is metabolism” (Schrödinger, 1944). According to this view, the ultimate equilibrium is death, and thus survival depends on staying as far as possible from equilibrium.

These ideas of life as a non-equilibrium process have been very influential for describing molecular and cellular functions in systems biology, including adaptation, sensing and transportation (Gnesotto *et al.*, 2018; Lynn *et al.*, 2021). Recently, research has demonstrated that non-equilibrium thermodynamics is also very useful for describing macroscopic systems such as the human brain (Deco *et al.*, 2022; Kringelbach *et al.*, 2024a; Lynn *et al.*, 2021), which is important since the brain is perhaps the main driver of how organisms avoid equilibrium and death.

The brain performs computation to avoid equilibrium but a main unanswered question is how to quantify the energy cost of these non-equilibrium computations (Bullmore and Sporns, 2012; Quintela-López *et al.*, 2022). The quantification of energy consumption of the brain is not straightforward since energy demands vary over time depending on the complexity of problems that the brain is forced to solve to survive. Still, careful studies of glucose consumption (which increases linearly with neuronal spike frequency) have shown that even though the human brain is only taking up 2% of total body weight, it accounts for 20% of resting metabolism (Raichle, 2010). Of this overall energy budget, the majority of the brain’s ‘dark energy’ (Raichle, 2006) is used for ongoing spontaneous activity (Clarke and Sokoloff, 1999; Raichle, 2006; Raichle *et al.*, 2001), while there is only a relatively small increase (of up to 5%) of extra energy used for cognition (Jamadar *et al.*, 2025; Raichle, 2010). This is all happening on a very small energy budget, where it has been estimated that the brain continuously runs on around 20 watts of power (Balasubramanian, 2021; Levy and Calvert, 2021). In contrast, a typical large high-performance computing cluster use up to six orders of magnitude more power to operate at around 2 megawatts. A major challenge for neuroscience is to understand why the cost of brain cognition is so much more efficient in biological brains.

The brain’s energy budget is used to enable the brain to flexibly map input with output through internal transformations, sometimes defined as ‘computation’ (Wolpert *et al.*, 2024). In other words, computation allows the brain to flexibly change its internal dynamics according to the mapping needed by modifying the distribution over the variables of the time-evolving system (Landauer, 1961; Wolpert, 2019). Importantly, this brain computation is distributed among communicating brain networks (Deco *et al.*, 2025). Most of the cost of distributed computation is much larger than local computation as convincingly demonstrated by Levy and Calvert (Levy and Calvert, 2021). In fact, glucose metabolism has been shown to scale linearly with the degree of functional connectivity in individual brains (Castrillon et al., 2023).

However, complementary to such metabolic estimates, here we use the revolutionary progress in stochastic thermodynamics (Peliti and Pigolotti, 2021; Seifert, 2012; Shiraishi, 2023; Van den Broeck and Esposito, 2015; Wolpert *et al.*, 2024) together with the most recent advances in modelling of whole-brain activity (Deco and Kringelbach, in press; Kringelbach and Deco, 2020) to provide the necessary tools to directly expressing the close links between energy consumption, information processing and non-equilibrium thermodynamics.

The new discipline of stochastic thermodynamics provides the necessary tools for analysing heat, work and entropy at the level of individual trajectories generated in a non-equilibrium physical system (Peliti and Pigolotti, 2021; Seifert, 2012; Shiraishi, 2023; Van den Broeck and Esposito, 2015; Wolpert *et al.*, 2024). The brain is one such non-equilibrium system where the mechanistic interactions between constituent parts can be determined using whole-brain modelling. We developed a new COCO (COst of COgnition) framework to discover the real cost of non-equilibrium cognition in terms of the underlying computations. This framework provides the means for quantifying energy consumption and non-equilibrium levels not only across species but also over the time course of activity linked to cognitive tasks in humans.

Here, we report the results of using the COCO framework to do both. In order to do so, we leveraged the availability of neuroimaging data from different species to build whole-brain models of the data. This whole-brain modelling expresses the local neuronal dynamics using the Hopf model coupled through the anatomical brain connectivity (Deco *et al.*, 2017b). Previous research has shown that the best fit of this modelling is at the criticality of the local dynamics, which allows the linearisation of the Hopf model (Ponce-Alvarez and Deco, 2024). Crucially, applying the tools of stochastic thermodynamics together with this linearisation allow us to derive an analytical expression quantifying the explicit and mechanistic links between non-equilibrium, energy cost and information processing for a given brain state. We also show that this key global relationship holds at the level of individual brain regions used for the whole-brain modelling.

Using cross-species empirical data from functional resonance imaging of the resting state and structural connectivity obtained using diffusion imaging and tract tracing, the COCO framework found that the human brain uses significantly less energy than both non-human primates and mice.

Focusing directly on the energy cost of cognition in humans, we used large-scale human neuroimaging data of 970 healthy participants to show that the resting state uses significantly less energy that seven different cognitive tasks. Importantly, the COCO framework can quantify the exact amounts of energy consumption, non-equilibrium and information processing for each task over both time and space. The findings demonstrate the different time evolution of non-equilibrium and information processing, and hence computational energy cost, not only in different tasks but even for distinct phases of a given task.

Attesting to the power of the COCO framework, we show how the averages across time of information processing and energy are excellent complementary predictors for machine learning classification. The results also show that more energy is used for tasks requiring a more dispersed computation. Also, the results also showed that minimal energy consumption is always found at the optimal working point for a given task by causally modifying the model.

Overall, complementary to existing metabolic measurements, the COCO framework can directly quantify the energy cost of non-equilibrium computation from empirical neuroimaging data. We found that the human brain uses significantly less energy than other mammalian species. Moving beyond spontaneous activity, the COCO framework can also quantify the real cost of cognition in terms of the non-equilibrium and energetic demands for information processing over time and space. This provides a deeper understanding of how evolution has shaped the biological brain to compute in a highly energy efficient way, which may ultimately provide new insights into minimising the energy use of current generations of digital computers and artificial intelligence.

## Results

The current state-of-the-art for estimating the energy costs associated with the brain is provided by metabolic neuroimaging (Jamadar *et al.*, 2025). Yet, more fine-grained measures such the computational and information processing cost of brain activity are hard to estimate from such glucose consumption dependent measures. Complementary, recent advances in stochastic thermodynamics has shown great promise for provide deeper insights into the thermodynamics of mind (Kringelbach *et al.*, 2024a), given that computation can be understood as the process allowing the brain to change the distribution over the underlying variables of the time-evolving system efficiently defining a map between input and output (**Figure 1A**).

**Figure 1.**
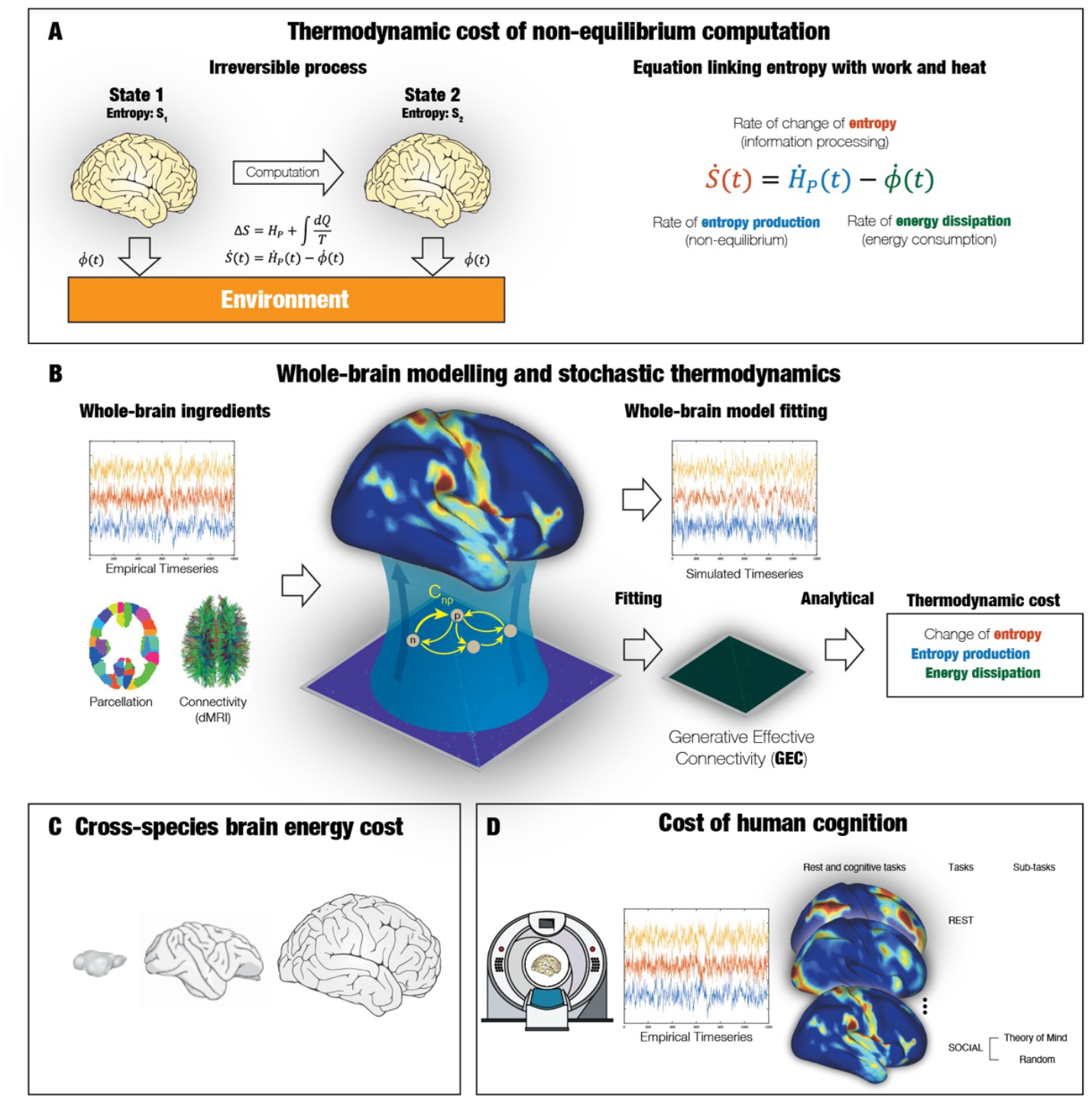
Framework for thermodynamic cost of non-equilibrium computation. **A)** The tools of stochastic thermodynamics can be used to compute three quantities, namely non-equilibrium level using entropy production, energy measured as the entropy flow exchanged between the brain and environment and information processing given by the change of entropy in the computing brain. The left panel of the figure shows a sketch of these entities, while the right panel shows the central equation linking the rate of change of information processing (entropy) with the rates of change of entropy production (work) and energy dissipation (heat). **B)** Whole-brain modelling is needed for this framework. The figure shows the ingredients needed, namely the structural connectivity and time series in a given parcellation. For the local dynamics the whole-brain model uses a linearised Hopf equation in each region to fit the data by optimising the GEC (generative effective connectivity) for each individual. This whole-brain model (with its individualised GEC) allows for an analytical expression quantifying the information processing, non-equilibrium and energy consumption at the global and regional levels (see Methods). **C)** Estimating the energy consumption across species, this framework is used on similar neuroimaging resting state data from human, non-human primates and mice. **D)** In order to further quantify the actual cost of cognition, the framework is used on the large-scale Human Connectome human neuroimaging dataset with seven different tasks and resting state. This estimation of the cost of human cognition establishes a direct relationship between energy consumption and computation in non-equilibrium states.

In the brain, computation can be defined as the change in the internal entropy of the system. Stochastic thermodynamics establishes that this change in brain entropy is related with the level of non-equilibrium and the energy dissipated to the environment. The level of non-equilibrium is described by the flux of entropy between the brain and the environment. As shown in the right panel of **Figure 1A** and in the *Methods*, the key equation expresses the relationship between rate of change in information processing (entropy) in terms of the rates of change of entropy production (work) and energy dissipation (heat).

**Figure 1B** shows that to quantify brain computation, the further ingredient of whole-brain modelling is needed. The whole-brain model is constructed from empirical structural connectivity and time series and can generate the empirical functional neuroimaging data (see full mathematical details in the *Methods*). Crucially, in order to obtain an analytical expression for quantifying the information processing, non-equilibrium and energy consumption at the global and regional levels, the local dynamics are modelled using a linearised Hopf model in each region to fit the data by optimising the GEC (generative effective connectivity) for each individual (see *Methods*).

Here, we use the COCO framework to estimate the energy consumption across species (**Figure 1C**) and to quantify the actual cost of cognition in large-scale neuroimaging data of seven different tasks and resting state (**Figure 1D**).

### Energetic computational costs across mammalian species

We first applied COCO to discover the computational costs across mammalian species (humans, non-human primates and mice) using the empirical data used in individual whole-brain models combining the time series from resting state functional magnetic resonance imaging (fMRI) with species-specific structural connectivity (collected using diffusion tractography in humans, DWI and tract-tracing in non-human primates and tract-tracing in mice). We used empirical data from 100 human individuals in two different parcellations (DK62 (Klein and Tourville, 2012) and Schaefer100 (Schaefer *et al.*, 2018)), 10 non-human primates in the COCOMAC82 parcellation (Kötter and Wanke, 2005) and 10 mice in the mouse 72 parcellation (Fulcher *et al.*, 2019).

The COCO framework estimated the generative effective connectivity (GEC) for each individual whole-brain model, which allows for the estimation of the energy consumption for each individual as the integral in time of the rate of the entropy flow (as specified in **Equation 25**). **Figure 2** shows that the lowest energy consumption was found in humans compared to both non-human primates and mice (p<0.001, Wilcoxon), while the highest energy consumption was found in mice.

**Figure 2.**
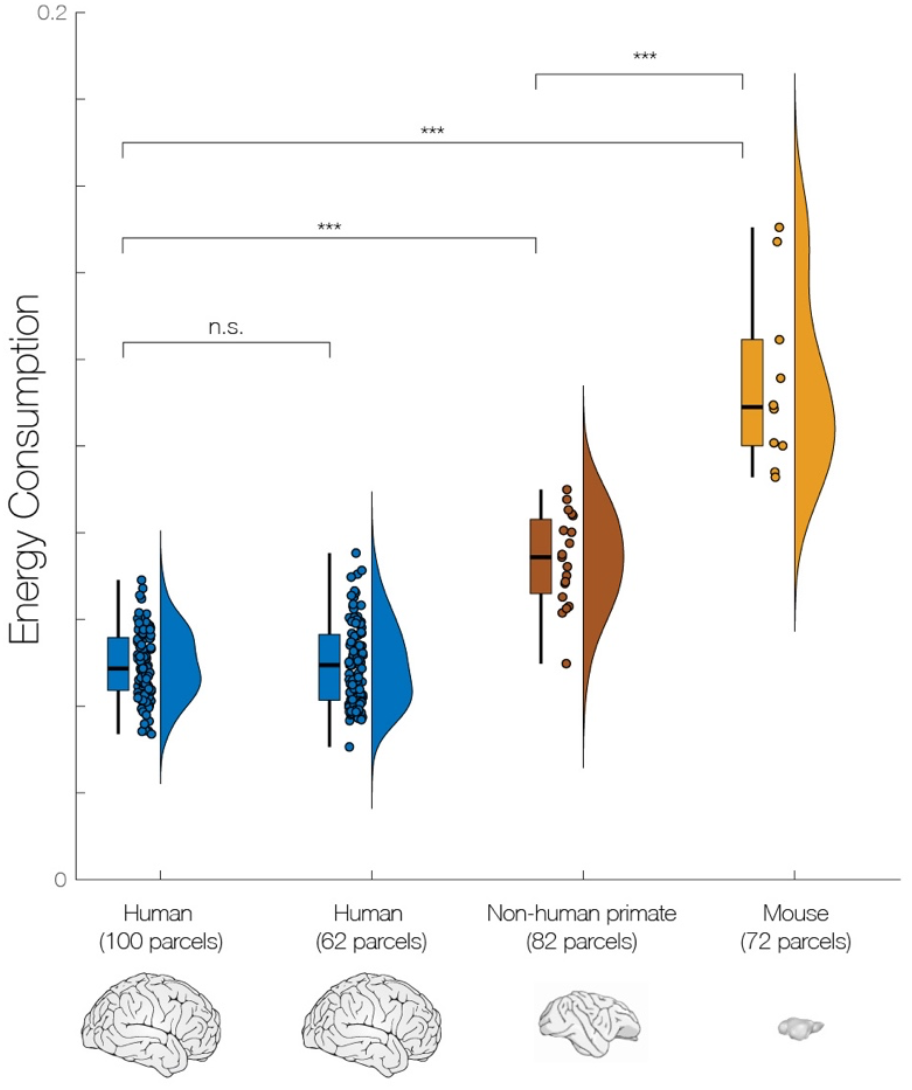
Energy consumption across different mammalian species. Whole-brain models were used to model time series from resting state functional magnetic resonance imaging (fMRI) collected in humans, non-human primates and mice. This was coupled with the species-specific structural connectivity collected using diffusion tractography in humans, DWI and tract-tracing in non-human primates and tract-tracing in mice. The results show significantly less energy consumption in humans compared to both non-human primates and mice. The energy consumption was highest in mice, suggesting an evolutionary optimisation of the effectiveness of computation.

### Cost of human cognition

Beyond the overall differences in resting state computation across species, the COCO framework uniquely allows for direct spatiotemporal estimation of the computational costs of any cognitive task by estimating both the global and regional thermodynamics measures.

For this we used the state-of-the-art Human Connectome Project (HCP) with neuroimaging data from 970 participants who in addition to resting state, all performed the standard battery of seven cognitive tasks, covering a broad range of human abilities in several major domains sampling the diversity of the brain (Barch *et al.*, 2013).

Importantly, the COCO framework allows for direct quantification of the exact regional levels of energy consumption, non-equilibrium and information processing for each task over time and space (where the regional levels are given by **Equation 34**). **Figure 3A** shows that the computational costs as measured by the global information processing is significantly different between rest and all seven tasks (integration over time of the global level given by **Equation 23**) (p<0.001, Wilcoxon).

**Figure 3.**
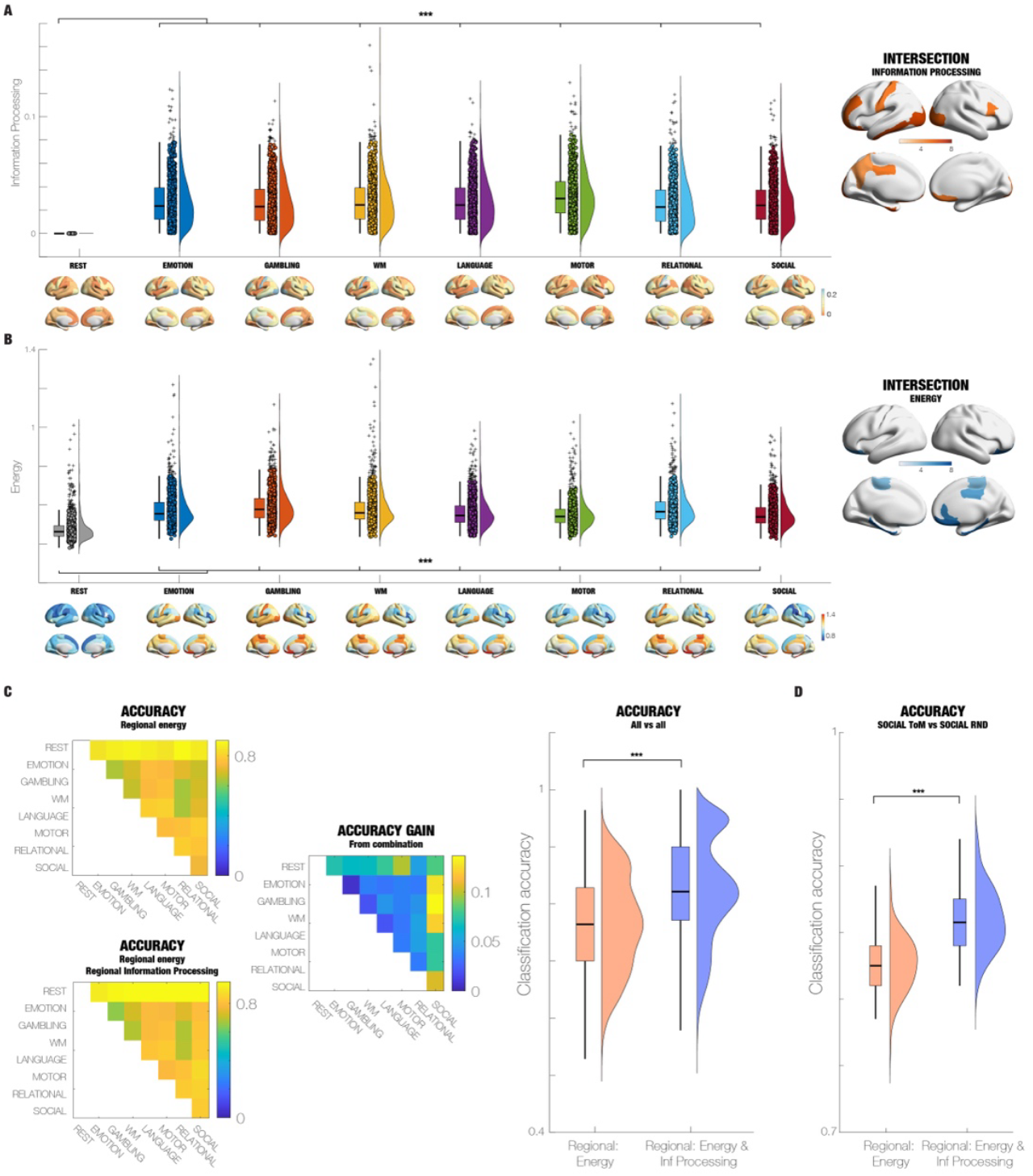
Energy consumption and information processing provide complementary information for understanding cognition. **A)** Information processing at the global level is significantly different between rest and all seven tasks (p<0.001, Wilcoxon). Equally, as can be seen from the renderings of the regional level of information processing (shown below rest and tasks), this is differently dispersed across space. The intersection across the seven tasks of the top regions for information processing (within the 20% quantile) is shown rendered on the right panel (where the darkness of orange indicates the number of tasks with a shared top region). One of the top region for information processing is the orbitofrontal cortex, intersecting across all 7 tasks. **B)** This panel shows the same information but now for global level of energy consumption, which is significantly different between rest and all seven tasks (p<0.001, Wilcoxon). Again, one top region for energy consumption is the orbitofrontal cortex, intersecting across the 7 tasks. **C)** In order to show the significantly dispersions of regional information processing and energy consumption, these regions across 970 participants were used for classification between all conditions (all pairwise combinations of rest and tasks). The resulting 8×8 matrix shows the highly significant ability to classify using a validation dataset with 1000-folds for using regional energy (top matrix) and regional energy and regional information processing (bottom matrix). As demonstrated by the difference in accuracy scores (middle matrix), using both measures provides more information than using just one of them, demonstrating the content of information of these thermodynamic regional measurements and their complementarity. This is also demonstrated by the violin plot of the accuracy for all pairwise combinations (over all 1000-folds), shown on the right of the panel. **D)** Even for more difficult distinctions of cognitive processing such as distinguishing between different phases of a task, these regional measures were highly predictive and complementary. This is demonstrated by the violin plot of accuracy of distinguishing Theory of Mind (ToM) vs random phases of the social task.

Interestingly, each task is characterised by different regional distributions across space as can be seen from the renderings of regional information processing in rest and seven tasks. Unlike the global levels of information processing, this regional information is sufficient for distinguishing the different tasks from each other (**Figure 3A**, upper panel).

Some brain regions are always in the top 20% quantile of regions for information processing, which can be see from the intersection of these regions rendered on the right side of the panel of **Figure 3A**. Here the darkness of orange indicates the number of tasks with a shared top region. The top regions for information processing found in all seven tasks are left entorhinal, left inferior temporal, left lateral occipital and left lateral orbitofrontal cortices. The top regions found across six tasks are left rostral middle frontal and right lateral occipital cortices, while top regions across five tasks are prefrontal regions (right pars opercularis and right medial orbitofrontal cortices), left postcentral and left posterior cingulate cortices. Finally, the left precuneus is found across four tasks.

Equally, the COCO framework can be used to quantify the energy consumption at the global level (**Equation 25**, integrated in time). Again, this shows that energy consumption in rest is significantly different from that used in all seven tasks (p<0.001, Wilcoxon). However, the global level of energy consumption does not distinguish the individual tasks. Instead, using the dispersed regional levels (using **Equation 36**, integrated in time) for each task distinguish the different tasks from each other (**Figure 3B**, lower panel).

In terms of the intersection of the top 20% quantile of regions for energy consumption, the rendering on the far right of **Figure 3B** shows these with the darkness of blue indicating the number of tasks with a shared top region. The top regions for information processing found in all seven tasks are the bilateral entorhinal and parahippocampal regions as well as prefrontal regions (right medial orbitofrontal and lateral orbitofrontal cortices). Top regions across six tasks are left lateral orbitofrontal and right rostral anterior cingulate cortices, while top regions across five tasks are right posterior cingulate cortex. Finally, the bilateral paracentral cortex is found across four tasks.

**Figure 3C** shows the SVM classification of all to all eight conditions (rest and seven tasks) using regional information processing (top) and the combination of regional energy consumption and information processing (bottom). In both cases, the resulting 8×8 matrix shows the highly significant ability to classify using a validation dataset with 1000-folds.

More interestingly, these measures are complementary since the difference in accuracy scores (middle matrix) are highly significant. Using both measures provides more information than using just one of them, which demonstrates the content of information of these thermodynamic regional measurements and their complementarity. Another way of showing this is provided by the violin plot of the accuracy for all pairwise combinations (over all 1000-folds), shown on the right of the panel (p<0.001, Wilcoxon).

**Figure 3D** shows that for even far more difficult distinctions between different phases of a task, the regional measures provided by the COCO framework were highly predictive and complementary. The violin plot shows the accuracy for distinguishing Theory of Mind (ToM) from random phases of the SOCIAL task (p<0.001, Wilcoxon).

### Dispersion of information processing is important for the energetic cost of cognition

Given that the metabolic cost of distributed computation has been shown to be much larger than local computation (Levy and Calvert, 2021), this raises the question of the computational cost for distributed computation. The COCO framework provides a direct answer to this question, given that the key analytical expression derived from combining thermodynamics with whole-brain modelling can be used both at the global as well as the regional level (see Methods). We were interested in studying the effects of distributed computation by analysing the dispersion of information processing across all regions in the whole brain.

**Figure 4A** shows a beautiful, highly significant monotonic correlation between the global energy consumption (mean across brain regions, using **Equation 36**) and the spatial dispersion of information processing (Dispersion IP, standard deviation across brain regions, using **Equation 34**) for all the seven cognitive tasks (across 970 participants) (r=0.99, p<0.001, Wilcoxon). This strongly suggests significantly different distributed computation for different tasks, which each require different mean levels of energy consumption.

**Figure 4.**
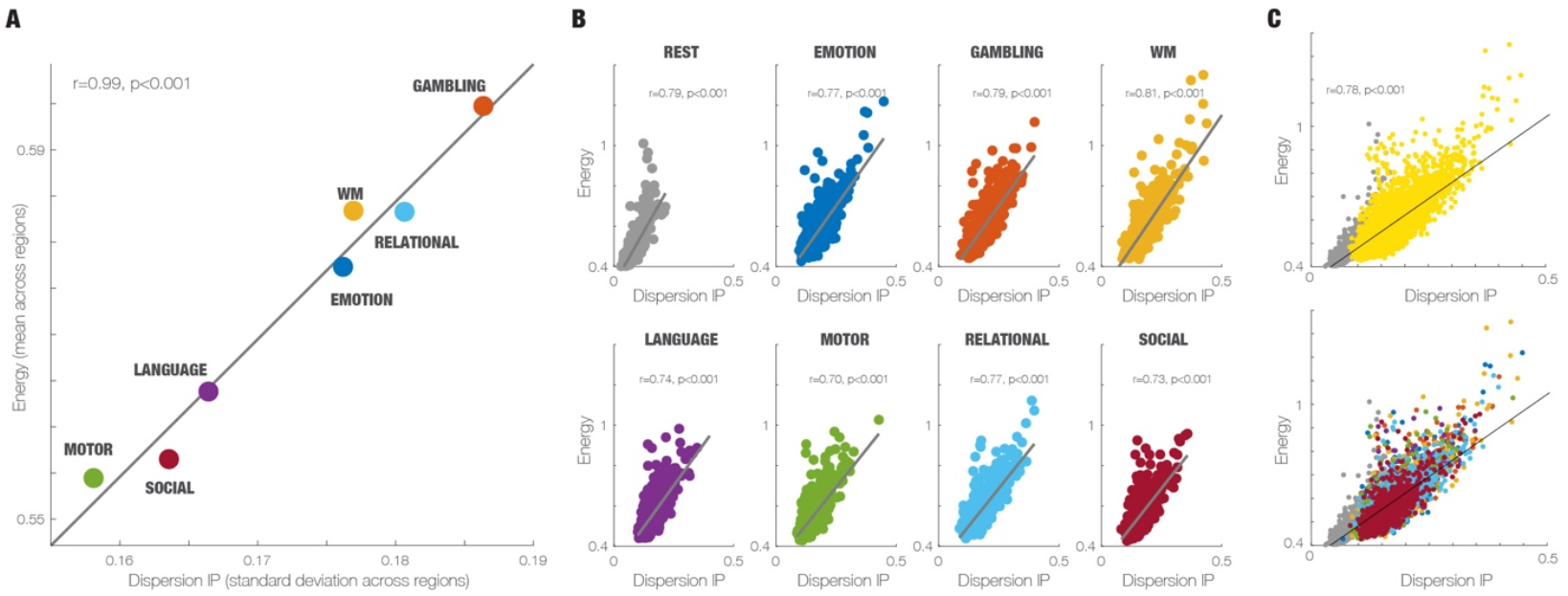
Cost of cognition is a function of the dispersion of information processing. **A)** The scatterplot shows a highly significant monotonic relationship between the global energy consumption (mean across brain regions) and the spatial dispersion of information processing (Dispersion IP, standard deviation across brain regions) for all seven cognitive tasks (across 970 participants). **B)** Exploring this further, the eight panels show the global energy consumption as a function of Dispersion IP for rest and the seven cognitive tasks. Here each dot represents the results one of the 970 participants. As can be seen, all correlations are highly significant (p<0.001, Wilcoxon). C) Finally, the scatterplots combines the eight panels into two figures. The top panel shows how rest (grey dots) is different from all seven tasks (yellow dots). The bottom panel shows scatterplot for the tasks.

**Figure 4B** provides another way of showing the global energy consumption as a function of Dispersion IP for rest and each of the seven cognitive tasks, where each point represents the results from one of the 970 participants. All eight correlations are highly significant (p<0.001 Wilcoxon). Similarly, **Figure 4C** combines the eight conditions into two figures, showing first shows how rest (grey dots) is different from all seven tasks (yellow dots).

### The cost of cognition is higher for tasks than when resting

Raichle has convincingly argued that most energy consumption in the brain is linked with what he has called ‘dark energy’, namely the energy associated with the brain’s spontaneous activity (Clarke and Sokoloff, 1999; Raichle, 2006; Raichle *et al.*, 2001). Research has shown that tasks only use a relatively small increase of up to 5% extra energy (Jamadar *et al.*, 2025; Raichle, 2010).

Beyond the global metabolic cost of cognition, this raises additional interesting question regarding the temporal evolution of energy consumption. The COCO framework can be used to quantify this and even show how this links with non-equilibirum and information processing.

Figure 5. shows how the COCO framework can used on the SOCIAL task which is an example of an engaging, reliable and validated task used for measuring social cognition (Castelli *et al.*, 2000). Participants are shown short video clips of twenty seconds of objects (circles, squares, triangles) that are either interacting in some way (suggestive of theory of mind, ToM), or moving randomly (RND). Each run has five blocks (three ToM and two RND) with five fixation blocks with a duration of 15 seconds.

As stated in the Methods, the thermodynamics measures are derived from the GEC associated with a given whole-brain model. We produced three different whole-brain models for each of the three conditions (ToM, RND and fixation cross). For each moment in time, we detect which GEC model is active, update **Equation 15** with the corresponding GEC (called ***J*** in the equation) and compute the three thermodynamic rate measurements of information processing, entropy production and energy consumption (from **Equations 23-25**, respectively, using the current ***J***). The absolute values of these three rate measures (averaged over participants) are shown in **Figure 5A**, overlaid on the three conditions. As can be clearly seen, these measures are clearly different between the ToM (in light red), RND (dark red) and fixation cross (no colour) conditions.

**Figure 5.**
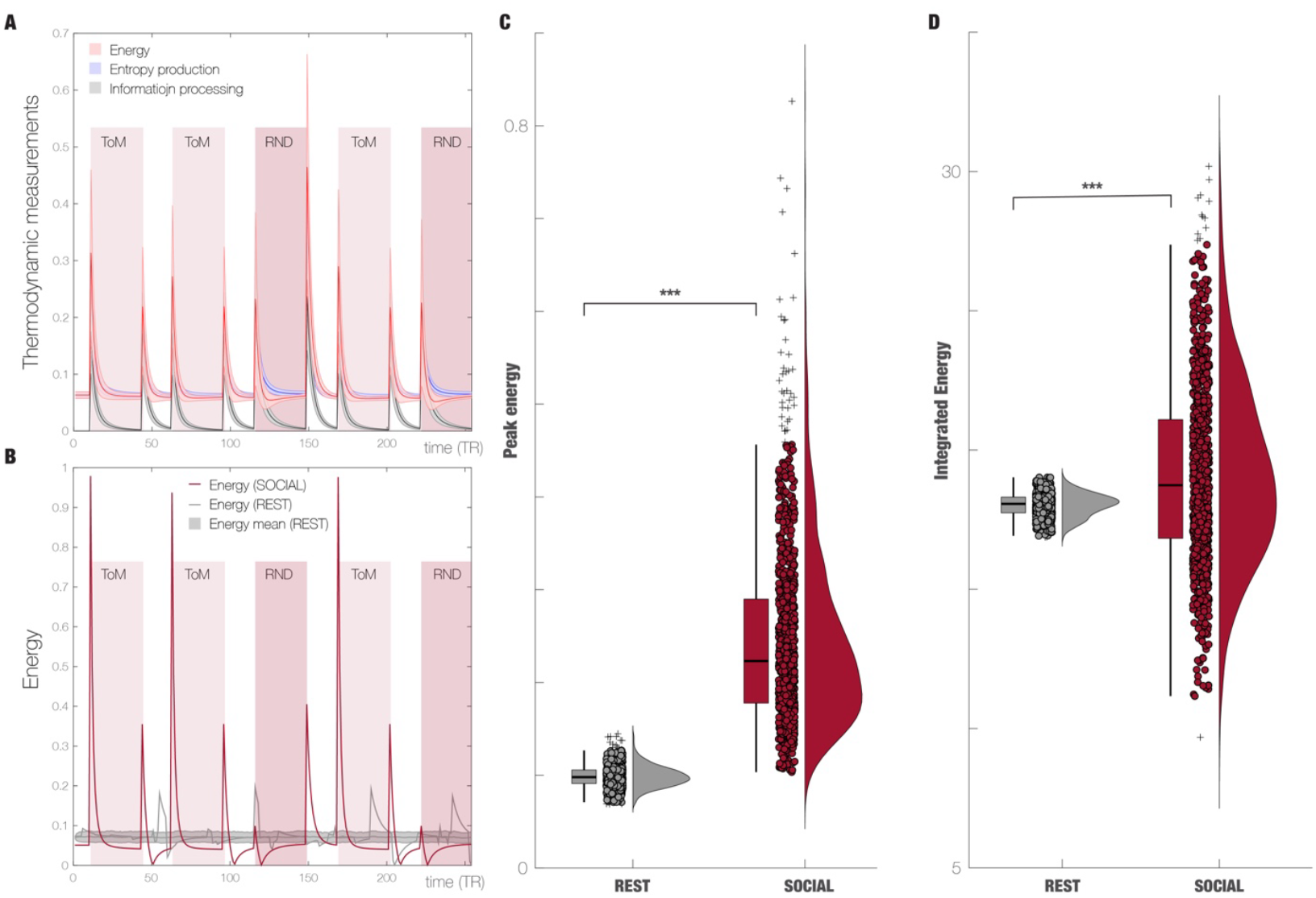
The cost of cognition is higher in task than when resting. **A)** The SOCIAL task involves making sense of interacting figures on a screen with two distinct phases of interpretation of either Theory of Mind (ToM, light red square) or random interactions (RND, darker red square). In between these phases there are periods of no active task engagement. The figure shows the average time evolution of the rate of change for energy (light red line), entropy production (light blue line) and information processing (grey line) across 970 participants and the standard deviations are indicated by the shadows. Clear peaks in the rate of change are seen for each of three measurements at the beginning of each phase, which reflect the need for resources for the necessary computations and the associated thermodynamic costs. The entropy production and information processing integrated over time for each phase are larger in RND than in ToM (not shown). **B)** Importantly, we compared the energy used for the task with the energy used in a separate resting state condition. The average energy consumption in the resting state with the mean energy (and standard deviation) across all participants is shown in grey. The energy used for the SOCIAL task for a single participant is overlaid on this (red lines). In addition, we show the energy used in the resting state, quantified by the novel Multicentred metastability method (see Methods). As can be seen, the peaks of energy consumption for the single individual are larger during task than resting. **C)** A full analysis of all participants showed a large significant increase of 150% of peak energy in the SOCIAL task compared to resting activity (p<0.001, Wilcoxon). **D)** When integrating over the total time, the total energy for SOCIAL vs resting is also significantly different (p<0.001, Wilcoxon) with an increase in the total energy of 5.4% commensurate with previous estimates of metabolic activity using PET (Raichle, 2010). Our thermodynamic results clearly demonstrate the cost of cognition, where the energy consumption for task is higher than for the resting state.

Directly comparing the evolution of the different conditions of the task with resting state requires a similar procedure for the resting state. For this, the COCO framework uses a similar method as used for task for resting state. However, given that resting state is not neatly composed into different conditions, this method uses clustering of the instantaneous phase-lock matrix of the resting state to establish seven models (e.g. seven models with different GECs). **Figure 5B** shows the energy consumption (red line) for a single participant aligned to the SOCIAL task as well as this participant’s resting state alignment (grey line). Obviously, participants will have different alignments of the resting state and therefore the peaks will be destroyed for the average energy consumption across all participants over time (grey area).

For the single participant, it would appear that the peak energy consumption is much larger for task than for resting state (compare red and grey lines). **Figure 5C** quantifies this observation to show that there is a significant increase of 150% of peak energy in the SOCIAL task compared to resting activity (p<0.001, Wilcoxon). This large increase would appear to contradict the much lower metabolic cost established using PET (Raichle, 2010). However, integrating over the total time, the total energy for SOCIAL vs resting shows a smaller increase in the total energy of 5.4% commensurate with previous estimates of metabolic activity (p<0.001, Wilcoxon).

## Discussion

Survival is one of the main goals of evolution (Schrödinger, 1944) and therefore the brain has been optimised to perform the necessary computation on a very limited energy budget. Here, we created a novel COCO (COst of Cognition) framework which provides a direct quantification of the energy consumption from neuroimaging data, based on recent progress in stochastic thermodynamics and whole-brain modelling. This allowed us to show that the cost of cognition is lowest in humans compared to non-human primates and mice. Equally, the global energy costs of human spontaneous resting activity are lower than when performing tasks.

Importantly, however, we were also able to quantify the energy cost on the regional level over the time course of an experiment. We showed that different kinds of tasks require distinct amounts of energy linked to distinct levels of information processing and non-equilibrium. These regional measurements of information processing and energy consumption are complementary and informative for describing a brain state as shown by our significant classification results of different tasks and subtasks.

The COCO framework uses the tools from stochastic thermodynamics (Peliti and Pigolotti, 2021; Seifert, 2012; Shiraishi, 2023; Van den Broeck and Esposito, 2015; Wolpert *et al.*, 2024). These are fundamental given that biological systems such as the brain are in non-equilibrium and therefore need non-equilibrium thermodynamics to describe the macroscopic features such as energy and information processing of a complex coupled system like the brain (Deco *et al.*, 2022; Kringelbach *et al.*, 2024a; Lynn *et al.*, 2021). Stochastic thermodynamics invariably uses modelling of systems to compute the underlying change in entropy as a function of entropy production and energy dissipation. Whole-brain modelling provides such a mechanistic model of empirical brain activity and allows for calculation of associated thermodynamic measures. In particular, the COCO framework uses the linearised Hopf equation to fit the empirical data (Deco *et al.*, 2017b; Ponce-Alvarez and Deco, 2024) and the resulting Langevin equation provide the necessary analytical spatiotemporal expression for thermodynamic measurements.

Central to cost of cognition is the cost of ‘computation’, which is a shapeshifting concept which is defined in many different ways. However, a useful definition of computation is as an operation that flexibly map input with output through internal transformations (Wolpert *et al.*, 2024). Inspired by the work of Wolpert and colleagues, the COCO framework is a potential solution for measuring the energetic cost of computation in the brain. Indeed, we applied stochastic thermodynamics together with whole-brain modelling to quantify the energy costs of the brain.

In accordance with previous findings of metabolic neuroimaging of the glucose consumption, the results show that the brain uses most of its energy budget to sustain the ‘dark energy’ used for ongoing spontaneous activity (Clarke and Sokoloff, 1999; Raichle, 2006; Raichle *et al.*, 2001). However, here the COCO framework also show that the peak energy is initially much larger (150%) when switching state within a task than the average much smaller change of 5.4% across the state. However, the average energy cost is very similar to that measured by metabolic neuroimaging (Jamadar *et al.*, 2025; Raichle, 2010).

Underlying the demand of energy for specific task computations, the brain uses many peaks of much lower energy to sustain the on-going spontaneous metastable activity (Hancock *et al.*, 2025), and to cover the rich repertoire of possible dynamics at the lowest energy cost. This is the reason why the average energy of resting dynamics (with many small peaks) is not so different from the average energy used in tasks, despite the large difference in task-related peaks of energy consumption found here.

Furthermore, these findings could provide an explanation of our previous results of how predominantly prefrontal regions are briefly recruited when performing tasks (Deco *et al.*, 2023; Kringelbach and Deco, 2024). This speculation is supported here by the prefrontal and in particular orbitofrontal regions using the most information processing and energy consumption across all tasks (Kringelbach, 2005).

Overall, the findings described here provide a direct quantification of cost of cognition, by way of calculating the energy cost of non-equilibrium computation from empirical neuroimaging data. The results demonstrate that the energy used by human brain is more efficient than for other mammalian species. The findings deepen our understanding of the highly energy efficient strategies that has evolved for computation in biological brains, which help with survival and even thriving (Kringelbach *et al.*, 2024b). The fundamental principles could potentially help develop far more efficient artificial computational devices.

## Methods

### Human Connectome Project: Acquisition and pre-processing

#### Ethics

The Human Connectome project is based on ethics obtained by the Washington University– University of Minnesota (WU-Minn HCP) Consortium who obtained full informed consent from all participants. Research procedures and ethical guidelines were followed in accordance with Washington University institutional review board approval (Mapping the Human Connectome: Structure, Function, and Heritability; IRB # 201204036).

#### Participants

We chose a sample of 970 participants, all of whom have resting state data and performed all seven tasks. These participants were selected from the March 2017 public data release from the Human Connectome Project (HCP).

#### The HCP task battery of seven tasks

All participants performed the standard HCP task battery described in detail on the HCP website (Barch *et al.*, 2013). This consists of seven tasks: Working memory (WM), motor, gambling, language, social, emotional, relational. According to HCP, the tasks were designed to cover a broad range of human abilities in several major domains sampling the diversity of the brain: 1) Visual, motion, somatosensory, and motor systems, 2) category-specific representations; 3) working memory, decision-making and cognitive control systems; 4) language processing; 5) relational processing; 6) emotion processing; and 7) social cognition. All HCP participants performed all tasks in two separate sessions (first session: gambling, WM and motor; second session: language, emotion processing, relational processing and social cognition).

#### 3T structural data

The HCP structural data were acquired using a customized 3 Tesla Siemens Connectom Skyra scanner with a standard Siemens 32-channel RF-receive head coil. For each participant, at least one 3D T1w MPRAGE image and one 3D T2w SPACE image were collected at 0.7 mm isotropic resolution.

#### Neuroimaging acquisition for fMRI HCP

All the 971 HCP participants were scanned on a 3-T connectome-Skyra scanner (Siemens). For the resting state data, we used one fMRI acquisition of approximately 15 minutes acquired on the first day, where participants had eyes open with relaxed fixation on a projected bright cross-hair on a dark background. We also used fMRI data from all the seven tasks. Full details of participants, the acquisition protocol and pre-processing of the data for both resting state and the seven tasks are provided by the HCP website (http://www.humanconnectome.org/) but here we give a brief summary. We used the standard minimal pre-processing of the HCP resting state and task datasets (as described in full details on the HCP website). This uses the standard HCP pipeline with standardized methods including FSL (FMRIB Software Library), FreeSurfer, and the Connectome Workbench software (Glasser *et al.*, 2013; Smith *et al.*, 2013). This pre-processing includes correction for spatial and gradient distortions and head motion, intensity normalization and bias field removal, registration to the T1 weighted structural image, transformation to the 2mm Montreal Neurological Institute (MNI) space, and using the FIX artefact removal procedure (Navarro Schroder *et al.*, 2015; Smith *et al.*, 2013). Head motion parameters were regressed out and structured artefacts were removed by

Independent Component Analysis followed by FMRIB’s ICA-based X-noiseifier (Griffanti *et al.*, 2014; Salimi-Khorshidi *et al.*, 2014).

The final pre-processed timeseries are in HCP CIFTI greyordinates standard space and available in the same standard surface-based CIFTI file for each participant for resting state and each of the seven tasks.

From these pre-processed timeseries, we extracted the average timeseries in two parcellations: Schaefer100 parcellation with a total of 100 cortical regions (50 regions per hemisphere) (Schaefer *et al.*, 2018) and DK62 with 62 cortical regions (31 regions per hemisphere) (Klein and Tourville, 2012). Using a custom-made Matlab script using the ft_read_cifti function from the Fieldtrip toolbox (Oostenveld *et al.*, 2011) to extract and average in each region. We filtered these average BOLD timeseries using a second-order Butterworth filter in the range of 0.008-0.08Hz.

### Non-human primate fMRI data

The non-human primate MRI data were made available as part of the Primate neuroimaging Data-Exchange (PRIME-DE) monkey MRI data sharing initiative, a recently introduced open resource for non-human primate imaging (Milham *et al.*, 2018).

#### Macaque dataset description

We used fMRI data from rhesus macaques (Macaca mulatta) scanned at Newcastle University. This samples includes 14 exemplars (12 male, 2 female); Age distribution: 3.9-13.14 years; Weight distribution: 7.2-18 kg (full sample description available online: *http://fcon_1000.projects.nitrc.org/indi/PRIME/files/newcastle.csv* and *http://fcon_1000.projects.nitrc.org/indi/PRIME/newcastle.html*).

Out of the 14 total animals present in the Newcastle sample, 10 had awake resting-state fMRI data; of these 10, all except the first had two scanning sessions available: to maximise our statistical power, these repeated sessions were included in the analysis. Thus, the total was 19 distinct sessions across 10 individual macaques, as in our previous publication (Luppi *et al.*, 2022).

#### Ethics approval

All of the animal procedures performed were approved by the UK Home Office and comply with the Animal Scientific Procedures Act (1986) on the care and use of animals in research and with the European Directive on the protection of animals used in research (2010/63/EU). We support the Animal Research Reporting of In Vivo Experiments (ARRIVE) principles on reporting animal research. All persons involved in this project were Home Office certified, and the work was strictly regulated by the U.K. Home Office. Local Animal Welfare Review Body (AWERB) approval was obtained. The 3Rs principles compliance and assessment was conducted by National Centre for 3Rs (NC3Rs). Animal in Sciences Committee (UK) approval was obtained as part of the Home Office Project License approval.

#### Animal care and housing

All animals were housed and cared for in a group-housed colony, and animals performed behavioural training on various tasks for auditory and visual neuroscience. No training took place prior to MRI scanning.

#### Macaque MRI acquisition

Animals were scanned in a vertical Bruker 4.7T primate dedicated scanner, with single channel or 4-8 channel parallel imaging coils used. No contrast agent was used. Optimization of the magnetic field prior to data acquisition was performed by means of 2nd order shim, Bruker and custom scanning sequence optimisation.

Animals were scanned upright, with MRI compatible head-post or non-invasive head immobilisation, and working on tasks or at rest (here, only resting-state scans were included). Eye tracking, video and audio monitoring were employed during scanning.

Resting-state scanning was performed for 21.6 minutes, with a TR of 2600ms, 17ms TE, Effective Echo Spacing of 0.63ms, voxels size 1.22 × 1.22 × 1.24. Phase Encoding Direction: Encoded in columns. Structural scans comprised a T1 structural, MDEFT sequence with the following parameters: TE: 6ms; TR: 750 ms; Inversion delay: 700ms; Number of slices: 22; In-plane field of view: 12.8 × 9.6cm2 on a grid of 256 × 192 voxels; Voxel resolution: 0.5 × 0.5 × 2mm; Number of segments: 8.

#### Macaque functional MRI preprocessing and denoising

The macaque MRI data were preprocessed using the *Pypreclin* pipeline for non-human primate MRI analysis *(https://github.com/neurospin/pypreclin)*, which addresses several specificities of monkey research. The pipeline is described in detail in the associated publication (Tasserie *et al.*, 2020). Briefly, it includes the following steps: (i) Slice-timing correction. (ii) Correction for the motion-induced, time-dependent B0 inhomogeneities. (iii) Reorientation from acquisition position to template; here, we used the recently developed National Institute of Mental Health Macaque Template (NMT): a high-resolution template of the average macaque brain generated from in vivo MRI of 31 rhesus macaques (Macaca mulatta) (Seidlitz *et al.*, 2018). (iv) Realignment to the middle volume using FSL MCFLIRT function. (v) Normalisation and masking using Joe’s Image Program (JIP)-align routine (*http://www.nmr.mgh.harvard.edu/~jbm/jip/*, Joe Mandeville, Massachusetts General Hospital, Harvard University, MA, USA), which is specifically designed for preclinical studies: the normalisation step aligns (affine) and warps (non-linear alignment using distortion field) the anatomical data into a generic template space. (vi) B1 field correction for low-frequency intensity non-uniformities present in the data. (vii) Coregistration of functional and anatomical images, using JIP-align to register the mean functional image (moving image) to the anatomical image (fixed image) by applying a rigid transformation. The anatomical brain mask was obtained by warping the template brain mask using the deformation field previously computed during the normalization step. Then, the functional images were aligned with the template space by composing the normalization and coregistration spatial transformations. Denoising of the macaque functional MRI data was carried out using the aCompCor denoising method implemented in the CONN toolbox. White matter and CSF masks were obtained from the corresponding probabilistic tissue maps of the high-resolution NMT template (eroded by one voxel); their first five principal components were regressed out of the functional data, as well as linear trends and 6 motion parameters (3 translations and 3 rotations) and their first derivatives. Finally, data were bandpass-filtered in the range of 0.008-0.09Hz, as in our previous work with these data (Luppi *et al.*, 2022). Macaque functional data were parcellated according to the COCOMAC 82-ROI “Regional Mapping” cortical atlas of Kötter and Wanke (Kötter and Wanke, 2005), nonlinearly registered to the NMT template used for preprocessing.

### Mouse FMRI data

The mouse fMRI data used here have been previously reported (Gutierrez-Barragan *et al.*, 2022) and for clarity and consistency of reporting, where possible we use the same wording as in the original publication.

#### Animals and ethics

In vivo experiments were conducted in accordance with the Italian law (DL 26/214, EU 63/2010, Ministero della Sanita, Roma) and with the National Institute of Health recommendations for the care and use of laboratory animals (Gutierrez-Barragan *et al.*, 2022). The animal research protocols for this study were reviewed and approved by the Italian Ministry of Health and the animal care committee of Istituto Italiano di Tecnologia (IIT). All surgeries were performed under anesthesia.

Adult (< 6 months old) male C57BL/6J mice were used throughout the study. Mice were group housed in a 12:12 hours light-dark cycle in individually ventilated cages with access to food and water ad libitum and with temperature maintained at 21 ± 1 degrees centigrade and humidity at 60 ± 10%. All the imaged mice were bred in the same vivarium and scanned with the same MRI scanner and imaging protocol employed for the awake scans (see below).

#### Experimental groups and datasets

A group of mice (n = 10, awake dataset) underwent head-post surgery, scanner habituation and fMRI image acquisitions (Gutierrez-Barragan *et al.*, 2022), where the full surgical, habituation and scanner protocol is described. These scan constitute the awake rsfMRI dataset used throughout this study.

#### MRI data acquisition

For awake scanning, the mouse was secured using an implanted headpost the custom-made MRI-compatible animal cradle and the body of the mouse was gently restrained (for details of the headpost implantation and habituation protocol, see the original publication (Gutierrez-Barragan *et al.*, 2022)). All scans were acquired at the IIT laboratory in Rovereto (Italy) on a 7.0 Tesla MRI scanner (Bruker Biospin, Ettlingen) with a BGA-9 gradient set, a 72 mm birdcage transmit coil, and a four-channel solenoid receive coil. Awake rsfMRI scans were acquired using a single-shot echo planar imaging (EPI) sequence with the following parameters: TR/TE=1000/15 ms, flip angle=60 degrees, matrix=100 × 100, FOV=2.3 × 2.3 cm, 18 coronal slices (voxel-size 230 × 230 × 600 mm), slice thickness=600 mm and 1920 time points, for a total time of 32 minutes.

#### Functional MRI preprocessing, denoising, and timeseries extraction

Preprocessing of fMRI images was carried out as described in previous work (Gutierrez-Barragan *et al.*, 2022). Briefly, the first 2 minutes of the time series were removed to account for thermal gradient equilibration. RsfMRI timeseries were then time despiked (3dDespike, AFNI), motion corrected (MCFLIRT, FSL), skull stripped (FAST, FSL) and spatially registered (ANTs registration suite) to an in-house mouse brain template with a spatial resolution of 0.23 × 0.23 × 0.6mm^3^. Denoising involved the regression of 25 nuisance parameters. These were: average cerebral spinal fluid signal plus 24 motion parameters determined from the 3 translation and rotation parameters estimated during motion correction, their temporal derivatives and corresponding squared regressors. No global signal regression was employed. In-scanner head motion was quantified via calculations of frame-wise displacement (FD). Average FD levels in awake conditions were comparable to those obtained in anesthetized animals (halothane) under artificial ventilation (p = 0.13, t-test) (Gutierrez-Barragan *et al.*, 2022). To rule out a contribution of residual head-motion, we further introduced frame-wise fMRI scrubbing (FD > 0.075 mm). The resulting time series were band-pass filtered (0.01-0.1 Hz band) and then spatially smoothed with a Gaussian kernel of 0.5 mm full width at half maximum. The timeseries were trimmed to ensure that the same number of timepoints were included for all animals, resulting in 1414 volumes per animal. Finally, data were parcellated into 72 cortical symmetric regions from the Allen Mouse Brain Atlas (CCFv3).

### Species-specific connectomes

#### Human structural connectome

In order to reconstruct a high-quality average structural connectivity (SC) matrix for constructing the whole-brain model (using the DK62 and Schaefer100 parcellations), we obtained multi-shell diffusion-weighted imaging data from 32 participants from the HCP database (scanned for approximately 89 minutes). The acquisition parameters are described in detail on the HCP website (Setsompop *et al.*, 2013). The connectivity was estimated using the method described by Horn and colleagues (Horn *et al.*, 2017).

In summary, the data was processed using a generalized q-sampling imaging algorithm implemented in DSI studio (http://dsi-studio.labsolver.org). A white-matter mask was produced from segmentation of the T2-weighted anatomical images, co-registered the images to the b0 image of the diffusion data using SPM12. For each of the 32 HCP participants, 200,000 fibres were sampled within the white-matter mask. Fibres were transformed into MNI space using Lead-DBS (Horn and Blankenburg, 2016). The methods used the algorithms for false-positive fibres shown to be optimal in recent open challenges (Maier-Hein *et al.*, 2017; Schilling *et al.*, 2019). Accordingly, the risk of false positive tractography was reduced by using the tracking method achieving the highest (92%) valid connection score among 96 methods submitted from 20 different research groups in a recent open competition (Maier-Hein *et al.*, 2017).

#### Macaque structural connectome

Anatomical (structural) connectivity data were derived from the recent macaque connectome (Shen *et al.*, 2019), which combines diffusion MRI tractrography with axonal tract-tracing studies from the CoCoMac database (Bakker *et al.*, 2012; Kötter and Wanke, 2005), representing the most complete representation of the macaque connectome currently available. Structural connectivity data are expressed as a matrix in which the 82 cortical regions of interest are displayed in x-axis and y-axis. Each cell of the matrix represents the strength of the anatomical connection between any pair of cortical areas. For consistency with the human structural connectome, a symmetrised connectome was used, as in our previous work (Luppi *et al.*, 2024).

#### Mouse structural connectome

For the mouse structural connectome, we used a parcellated version of the high-resolution mouse connectome from the Gozzi lab (Coletta *et al.*, 2020). The high-resolution mouse structural connectome was obtained in the following way. The present mouse structural connectome is based on “high-resolution models of the mouse brain connectome (100 μm^3^) previously released by Knox and colleagues (Knox *et al.*, 2019). The Knox connectome is based on 428 viral microinjection experiments in C57BL/6J male mice obtained from the Allen Mouse Brain Connectivity Atlas (*http://connectivity.brain-map.org/*. The connectome data were derived from imaging enhanced green fluorescent protein (eGFP)–labeled axonal projections that were then registered to the Allen Mouse Brain Atlas and aggregated according to a voxel-wise interpolation model (Knox *et al.*, 2019). Before constructing the SC matrix, Coletta and colleagues ensured symmetry along the right-left axis for all the major macrostructures of the mouse brain. To this purpose, they “flipped each macrostructure (isocortex, hippocampal formation, subcortical plate, pallidum, striatum, pons, medulla, midbrain, thalamus, hypothalamus, cerebellum, and olfactory bulb) along the sagittal midline (once for the right hemisphere and once for the left hemisphere) and took the intersection with the respective nonflipped macrostructure. This procedure resulted in the removal of a set of nonsymmetric voxel (total fraction, 8.6%), the vast majority of which reside in fringe white/gray matter or cerebrospinal fluid/gray matter interfaces. The removal of these nonsymmetric voxels did not substantially affect the network structure of the resampled connectome, as assessed with a spatial correlation analysis between the symmetrized and non-symmetrised right ipsilateral (i.e., squared) connectome”. Coletta and colleagues then filtered out fibre tracts and ventricular spaces, and estimated SC using a resampled version of the recently published voxel scale model of the mouse structural connectome (Knox *et al.*, 2019), in order to make the original matrix computationally tractable. Resampling of the Knox et al. connectome was carried out by aggregating neighbouring voxels according to a Voronoi diagram based on Euclidean distance between neighbouring voxels, preserving the intrinsic architectural foundation of the connectome while minimizing spatial blurring and boundary effects between ontogenically distinct neuroanatomical divisions of the mouse brain, or white/grey matter, and parenchymal/ventricular interfaces. Please see Coletta and colleagues for details of the Voronoi aggregation scheme. By averaging the connectivity profile of neighbouring voxels based on their relative spatial arrangement, this strategy has also the advantage of mitigating limitations related to the enforced smoothness of source space used by the original kernel interpolation used by Knox and colleagues (Knox *et al.*, 2019).

A whole-brain connectome was then built under the assumption of brain symmetry (Coletta *et al.*, 2020). Forty-four dangling nodes (i.e., nodes with no outgoing connectivity) were next removed from the resulting matrix, resulting in a final weighted and directed 15,314 × 15,314 matrix composed of 0.027-mm^3^ aggregate Voronoi voxels. The obtained Voronoi diagram made it possible to map the results back into the original 100-μm three-dimensional coordinate system of the Allen Institute mouse brain connectome [CCFv3]. For each pair of the 72 Allen Atlas cortical regions that we employed, their structural connectivity was obtained by averaging the connectivity of the respective constituent voxels.

To facilitate comparison of results between human and other species, the structural connectomes of macaque and mouse were symmetrised, to avoid imposing structural asymmetries and instead allow the model itself to determine the most suitable level of asymmetry (if any). Similarly, since the human and macaque structural connectomes are sparse, whereas the mouse connectome is provided as fully dense, the latter was thresholded to 50% density. However, we demonstrate that our results are not dependent on either of these methodological choices, as shown in the Supplementary Information.

### Whole-brain model

The whole-brain model is the same across species. For the human case the modelling uses the DK62 parcellation (Klein and Tourville, 2012), except for the cross species comparison where we also used the Schaefer100 (Schaefer *et al.*, 2018). For the non-human primates, we use the COCOMAC82 parcellation (Kötter and Wanke, 2005), and for the mice we use the Mouse72 parcellation (Coletta *et al.*, 2020; Fulcher *et al.*, 2019).

The local dynamics of each brain region is expressed by a Stuart-Landau oscillator (i.e., as the normal form of a supercritical Hopf bifurcation). The Hopf model has become a standard model for examining the shift from noisy to oscillatory dynamics (Kuznetsov, 1998), and they have been used to replicate key aspects of brain dynamics observed in electrophysiology (Freyer *et al.*, 2011; Freyer *et al.*, 2012), magnetoencephalography (Deco *et al.*, 2017a) and fMRI (Deco *et al.*, 2019; Kringelbach *et al.*, 2020).

Here, for a parcellation with *N* regions, the coupling the local dynamics of *N* Stuart-Landau oscillators via the connectivity matrix ***C***, defines the whole-brain dynamics as follows

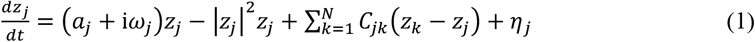

where for the oscillator in region *j*, the complex variable *z*_*j*_ denotes the state (*z*_*j*_ =*x*_*j*_ + i*y*_*j*_), *η*_*j*_ is additive uncorrelated Gaussian noise with variance σ^2^ (for all *j*), ω_*j*_ is the intrinsic node frequency, and *a*_*j*_ is the node’s bifurcation parameter. Within this model, the intrinsic frequency ω_*j*_ of each node is in the 0.008–0.08Hz band. The intrinsic frequencies were estimated from the data, as given by the averaged peak frequency of the narrowband blood-oxygen-level-dependent (BOLD) signals of each brain region. In **Equation 1**, *C*_*jk*_ is a specific entry in the coupling connectivity matrix ***C***, which is optimised to fit the resting-state empirical data, as detailed below. For *a*_*j*_ > 0, the local dynamics settle into a stable limit cycle, producing self-sustained oscillations with frequency ω_*j*_/(2***π)***. For *a*_*j*_ < 0, the local dynamics present a stable spiral point, producing damped or noisy oscillations in the absence or presence of noise, respectively. The fMRI signals were modelled by the real part of the state variables, i.e., *x*_*j*_ = Real(*z*_*j*_). It has been shown that the best working point for fitting whole-brain neuroimaging dynamics is at the brink of the bifurcation, i.e. with *a*_*j*_ slightly negative but very near to zero (usually *a*_*j*_ = −0.02) (Deco *et al.*, 2017b).

Importantly, the proximity to criticality is crucial, because it allows a linearization of the dynamics (Ponce-Alvarez and Deco, 2024), which permits an analytical solution for the functional connectivity matrix ***FC***^*model*^ (given by the Pearson correlations between all pairs of brain regions).

Briefly, we can estimate the functional correlations of the whole-brain network using a linear noise approximation (LNA). Hence, the dynamical system of *N* nodes (**Equation 1**) can be re-written in vector form as:

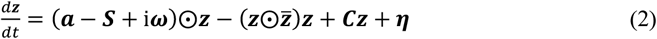

where ***z*** = [*z*_1_, …, *z*_*N*_]^*T*^, ***a*** = [*a*_1_, …, *a*_*N*_]^*T*^, ***ω*** = [ω_1_, …, ω_*N*_]^*T*^, ***η*** = [*η*_1_, …, *η*_*N*_]^*T*^ and ***S*** = [*S*_1_, …, *S*_*N*_]^*T*^ is a vector containing the strength of each node, i.e. *S*_*i*_ = ∑_*j*_ *C*_*ij*_. The superscript []^*T*^represents the transpose, 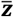 is the complex conjugate of ***z*** and ⨀ is the Hadamard element-wise product. As such, the equation describes the linear fluctuations around the fixed point ***z*** = 0, which is the solution of ^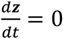^. Discarding the higher-order terms 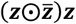 and separating the real and imaginary parts of the state variables, the evolution of the linear fluctuations follows a Langevin stochastic linear equation:

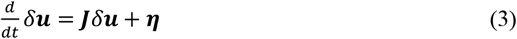

where the 2*N*-dimensional vector δ***u*** = [δ***x***, δ***y***]**^*T*^** = [δ*x*_1_, …, δ*x*_*N*_, δ*y*_1_, …, δ*y*_*N*_]^*T*^ contains the fluctuations of real and imaginary state variables. The 2*N* × 2*N* matrix ***J*** is the Jacobian of the system evaluated at the fixed point, which can be written as a block matrix

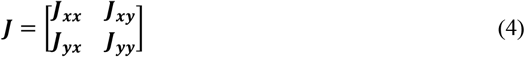

where ***J*_*xx*_, *J*_*xy*_, *J*_*yx*_, *J*_*yy*_** are *N* ×*N* matrices given as ***J*_*xx*_** = ***J*_*yy*_** = diag(***a*** − ***S***) + ***C*** and ***J*_*xy*_** = −***J*_*yx*_** = diag(***ω***), where diag(***v***) is the diagonal matrix whose diagonal is the vector ***v***. Please note that this linearisation is only valid if ***z*** = 0 is a stable solution of the system, that is if all eigenvalues of ***J*** have negative real parts. To compute the covariance matrix ***K*** = ⟨δ***u***δ***u*^*T*^**⟩, one can begin by writing **Equation 3** as *d*δ***u*** = ***J***δ***u****dt* +*d****W***, where *d****W*** is an 2*N*-dimensional Wiener process with covariance ⟨*d****W****d****W*^*T*^**⟩ = ***Q****dt*, where ***Q*** is the noise covariance matrix, which is diagonal if the noise is uncorrelated. Using Itô’s stochastic calculus, we get *d*(δ***u***δ***u*^*T*^**) =*d*(δ***u***)δ***u*^*T*^** + δ***u****d*(δ***u*^*T*^**) +*d*(δ***u***)*d*(δ***u*^*T*^**). Noting that ⟨δ***u****d****W*^*T*^**⟩ = 0, taking the expectations and keeping terms to first order in the differential *dt*, we obtain:

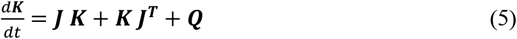

Hence, the stationary covariances (for which 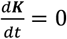) can be obtained by solving the following analytic equation:

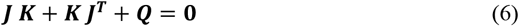

**Equation 6** is a Lyapunov equation that can be solved using the eigen-decomposition of the Jacobian matrix (Deco *et al.*, 2014). We obtained the simulated functional connectivity matrix ***FC***^*model*^ from the first *N* rows and columns of the covariance ***K***, which corresponds to the real part of the dynamics, and thus the BOLD fMRI signal.

### Optimizing the generative effective connectivity (GEC)

For defining the GEC of a whole-brain model, we first fit the model to the empirical neuroimaging data for each participant. For this, we used a noise *η*_*j*_ with variance σ^2^ = 0.02 homogeneous for all regions in a given parcellation. In order to create the generative effective connectivity (GEC), we used a pseudo-gradient procedure to optimise the initial coupling connectivity matrix ***C***, derived from the anatomical structural connectivity which was then iteratively used to create the GEC.

Specifically, we iteratively compared the output of the model with the empirical measures of the functional correlation matrix (***FC***^*empirical*^), i.e., the normalised covariance matrix of the functional neuroimaging data. Furthermore, we also compared the output of the model with the normalized ***τ***time-shifted covariances (***FS***^*empirical*^***(τ)***). These normalised time-shifted covariance matrices are generated by taking the shifted covariance matrix ***KS***^*empirical*^***(τ)*** and dividing each pair (*i, j*) by 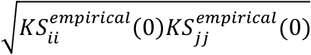.

Note that these normalised time-shifted covariances break the symmetry of the couplings and thus improve the level of fitting (Gilson *et al.*, 2017). Using a heuristic pseudo-gradient algorithm, we proceeded to update the ***C*** until the fit is fully optimised. More specifically, the updating uses the following form:

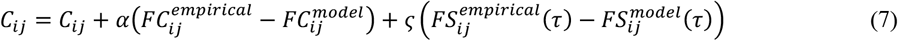

where 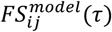 is defined similar to 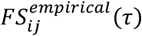. In other words, the updating is given by the first *N*rows and columns of the simulated ***τ***time-shifted covariances ***KS***^*model*^(***τ)*** normalised by dividing each pair (*i, j*) by 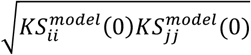, where ***KS***^*model*^(***τ)*** is the shifted generative covariance matrix computed as follows:

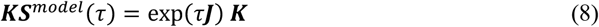

Please note that ***KS***^*model*^(0) = ***K***. The model was then run repeatedly with the updated ***C*** until the fit converges towards a stable value. As stated above, we initialised ***C*** using the average anatomical connectivity (see *Methods*above) and only update known existing connections from this matrix (in either hemisphere). There is one exception, however, to this rule which is that the algorithm also updates homologue connections between the same regions in either hemisphere, given that tractography is known to be less accurate when accounting for this connectivity.

For the Stuart-Landau model, we used ***α***= ***ς***= 0.000***4***and continue until the algorithm converges. For each iteration we compute the model results as the average over as many simulations as there are participants. In the human case, we applied this procedure to fit a whole-brain model to resting state data for each individual which generated 970 individual GECs, which were subsequently used in the framework described as follows. For non-human primates, we applied this procedure to 19 datasets and for the mice to 10 datasets.

### Analytical derivation of entropy production, energy dissipation and information processing

Non-equilibrium systems are central to many scientific fields, but fundamental challenges remain, especially in defining microscopic entropy production—a key measure of irreversibility. In thermodynamic terms, the change of entropy ***ΔS*** for a reversible process is given by:

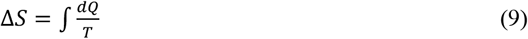

where *d****Q***is the heat exchanged and *T*is temperature. For irreversible processes,

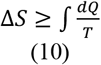

and the difference defines entropy production ***HP,*** i.e.:

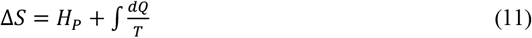

The rates of change derived with respect to time (with the dot on the top of a function signifying the time derivative) are given by:

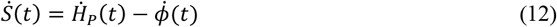

In this equation 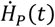 is the entropy production rate of the system, which (according to the second law of thermodynamics) is always non-negative, and 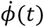 is the entropy flux rate from the system to the environment. For systems in a non-equilibrium steady-state, 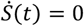which implies 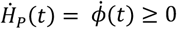. Please note that it is only for thermodynamic equilibrium that the equality in this equation is expected to hold.

***I***n the brain, 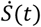 the rate of change of entropy, can be interpreted as ‘computation’, that is the flexible mapping of input with output through internal transformations and consequently as a measure of information processing. On the other hand, the 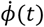 is the entropy flux can be interpreted as the energy consumption for performing a particular computation, and 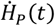 is the level of non-equilibrium. These fundamental thermodynamic quantities can be calculated analytically when the system is a linear Langevin system, which is exactly the case for our linearised Hopf model (described above in **Equation 3**).

Let us by consider following Langevin system using a simplifying notation of δ***u*= *x***:

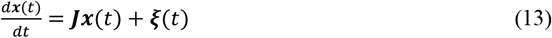

The ***ξ(****t*) are 2N independent standard Wiener processes with covariance:

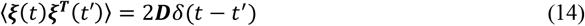

Therefore ***D***is a 2*N*× 2*N*diagonal matrix (but now generalised to the possibility that the diagonal elements could be different). The angular brackets indicate the mathematical expectation over time and the δ(*t*) is the Dirac***’s*** delta function.

Given this, it is straightforward to derive the temporal evolution of the covariance matrix, ***θ***= ⟨***x***(*t*)***x*^*T*^**(*t****′)***⟩ − ⟨***x***(*t*)⟩⟨***x***(*t****′)***⟩, which is given by:

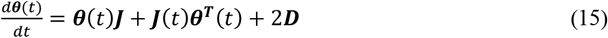

The probability distribution ***P(x***,*t*) associated with the Langevin equation is described by following Fokker-Planck equation:

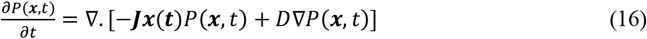

Where ***∇***denotes the spatial derivative with respect to ***x***. The above Fokker-Planck equation can be expressed as the following continuity equation:

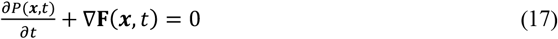

where the probability current is given by:

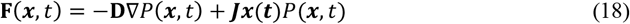

The stationary solution can be derived analytically and is given by following multivariate Gaussian distribution:

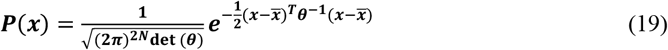

The entropy is defined by:

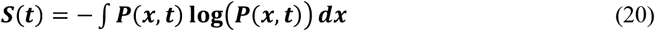

As shown by Landi and colleagues (Landi *et al.*, 2013) and Gilson and colleagues (Gilson *et al.*, 2023), the rate of change of entropy can be decomposed as:

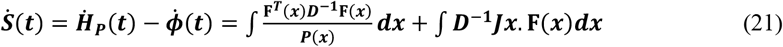

Where the stationary probability current ***F(x***) is given by

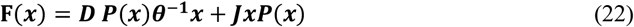

From these equations, following the lead of Landi and colleagues, we can express the information processing (as the rate of the change in entropy):

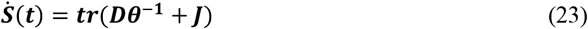

***w***hile the entropy production can be expressed as:

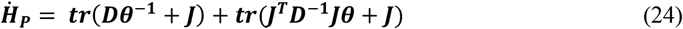

***a***nd energy consumption (measure as entropy flow out of the system) as follows:

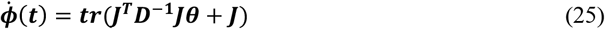

where *t****r***is the trace of the matrix.

### Linking symmetry with irreversibility

If the coupling matrix ***J*** is symmetric and the noise is the same in every direction, i.e., 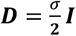, then the stationary equation for the covariance reads:

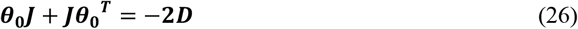

Since ***J*** is symmetric ***J*** = ***J*^*T*^** and thus

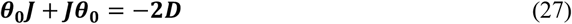

and consequently

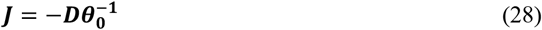

and thus

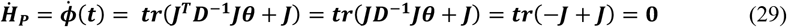

In summary, when ***J*** is symmetric and the noise is isotropic, the entropy production at steady state vanishes, and thus the system is in equilibrium. On the contrary, if ***J*** is asymmetric, the system is in non-equilibrium.

### Regional decomposition of the thermodynamic measurements

One can decompose the information processing, entropy production and energy consumption, in terms that involve only the interactions associated with each single node. Let us define an eigenvector decomposition of the covariance matrix:

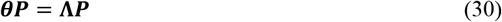

Where ***P***is the matrix with eigenvectors in each column, and ***Λ***the eigenvalues diagonal matrix. Let us define

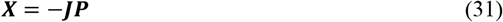

and

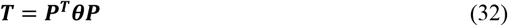

Using traces properties

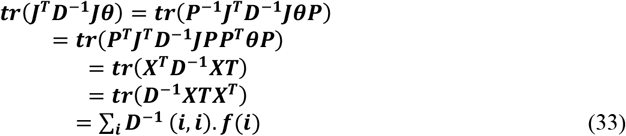

This decomposes the trace in a sum of terms, where in each term appears only elements of ***J*** associated with one node. Similarly, we can express all thermodynamic quantities for each node *i*so that information processing can be expressed by:

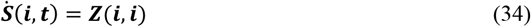

entropy production:

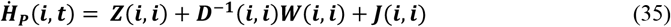

and energy consumption:

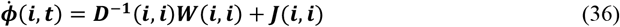

where

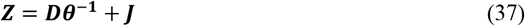

and

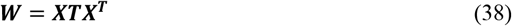

### SVM for pattern separation and classification

Both pattern separation and classification were conducted using a support vector machine (SVM) with linear kernels as implemented by the function *fitcecoc*in Matlab2024b, returning a full, trained, two class, error-correcting output codes model with the predictors in the input with class labels.

The input features used for classification were regional values for 1) information processing or 2) information and energy consumption. The SVM was trained using the leave-one-out cross-validation procedure, that is we randomly chose one participant for generalisation and used the whole rest for training. This was repeated and shuffled 1,000 times. Furthermore, the training set was balanced in terms of number of examples for each class label and randomly selecting the participant in each class for each shuffling iteration.

